# The Ferrous Iron Transporter FeoB1 is Essential for *Clostridioides difficile* Toxin Production and Pathogenesis in Mice

**DOI:** 10.1101/2022.03.03.482942

**Authors:** Aditi Deshpande, Chetna Dureja, Ann M. Mckelvey, Wei Zhao, Jacob T. Rutherford, Abiola O. Olaitan, Julian G. Hurdle

## Abstract

Ferrous iron is the dominant form of iron in the oxygen-limited large intestine, the site for *Clostridioides difficile* infection (CDI). We investigated the extent to which *C. difficile* requires the ferrous iron transporter 1 (FeoB1), a main permease for iron acquisition, using *in vitro* and *in vivo* approaches. Construction of *feoB1* deletion mutant in *C. difficile* R20291 (i.e., R20291Δ*feoB1*) decreased intracellular iron content by ∼25%. This was accompanied by downregulation of *tcdA* and *tcdB* genes and reduced synthesis of TcdA and TcdB toxins, which was reflected by a ∼1000-fold reduction in cytopathy against Vero cells. Complementation with WT *feoB1* restored toxin production. Transcriptional and biochemical analyses revealed Δ*feoB1* altered metabolic pathways that negatively impact toxin production, including downregulation of the oxidative branch of the Kreb cycle and an associated cellular accumulation of pyruvate. Hence FeoB1 influences multiple bioenergetic and redox pathways, which in turn inhibits toxin biosynthesis. R20291Δ*feoB1* was avirulent in mice with underlying colitis. In this model, mice are prone to develop severe CDI, but mice infected with the mutant lacked diarrheal symptoms, had lesser inflammatory responses, bacterial bioburdens and cecal toxin titers. These results strongly imply that ferrous iron acquisition and homeostasis is central to toxin metabolism and virulence. Hence targeting FeoB1 could be a potential therapeutic strategy to impede the colonization and pathogenesis of *C. difficile*, even in the setting of colitis where intestinal bleeding may occur and CDI is more severe.

## INTRODUCTION

*Clostridioides difficile* infection (CDI) is a leading cause of hospital-acquired diarrhea, with 30,600 and 20,500 in-hospital deaths in 2011 and 2017, respectively (1). The main risk factor for CDI is prior antibiotic use, which causes dysbiosis (2). Treatment with anti-*C. difficile* antibiotics, vancomycin, metronidazole or fidaxomicin, are associated with recurrence rates that may be 20% or higher (3–5). CDI symptoms range from mild-to-moderate diarrhea to severe pseudomembranous colitis, resulting from epithelial damage by *C. difficile* toxins TcdA and TcdB (6). These toxins glycosylate GTPases (e.g., Rho, Rac, Cdc42) to block actin polymerization, leading to cytoskeletal disruption, loss of barrier integrity, and ultimately epithelial and sub-epithelial cell death, which drives inflammation and tissue damage (6). While neutralizing these toxins is a validated clinical approach (7), targeting their biosynthesis represents a promising and emerging therapeutic strategy (8).

TcdA and TcdB are chromosomally encoded within the pathogenicity locus (PaLoc) and are subject to various levels of metabolic regulatory control (6). PaLoc encodes the following: alternative sigma factor TcdR, which activates transcription of *tcdA* and *tcdB*; the TcdR antagonist TcdC that represses *tcdA* and *tcdB*; and the holin-like protein TcdE that exports the toxins. TcdA and TcdB are produced in the late logarithmic and stationary phases of growth and are regulated by various environmental and nutritional changes. For example, *tcdA* and *tcdB* are repressed by metabolizable sugars (e.g., glucose) and metabolism of amino acids like proline, cysteine, and branched-chain amino acids (BCAAs) (9–11). Glucose activates catabolite repression through the global regulator CcpA that binds upstream of promoters in PaLoc, particularly the *tcdR* promoter, to block transcription of *tcdA* and *tcdB* (9). During logarithmic growth, high intracellular concentrations of GTP bind to the nutritional regulator CodY, which in turn binds to regions of the *tcdR* promoter (12). Pyruvate build-up also suppresses toxin production via an unresolved mechanism that might involve nutrient sensing and metabolic control via the YpdA/YpdB pyruvate sensing two-component system (11); pyruvate is pivotal to several anaplerotic reactions, linking glycolysis, amino acid metabolism, and the TCA cycle, and contributes to maintaining NAD^+^/NADH balance (13). Conversely, butyrate enhances the synthesis of toxins by an unknown mechanism that may be related to NAD+ regeneration via alternative fermentative pathways that produce butyrate (14, 15). These and other examples, all point to there being a complex metabolic and regulatory network for *C. difficile* toxin production.

Within environments that are anaerobic and acidic, such as the gastrointestinal tract, iron mainly exists as ferrous iron (16, 17). Gut bacteria must therefore be able to obtain ferrous iron in the microaerophilic and anaerobic niches of the intestine. *C. difficile* is thought to be limited in its iron capture mechanisms, as it appears to lack heme degrading enzymes related to heme oxygenases or staphylococcal IsdG (18, 19); it is also unable to robustly grow on transferrin or lactoferrin (20). *C. difficile* also encodes a ferric (Fe^3+^) hydroxamate transporter (FhuDBGC) to capture xenosiderophores, but its deletion did not attenuate colonization and virulence in mice (21). However, *C. difficile* stores ferrous iron (Fe²⁺) in structures known as ferrosomes that are made by FezA and FezB and deletion of FezA attenuates infectivity in mice (22). Thus, the acquisition of Fe²⁺ is crucial for *C. difficile* metabolism and virulence, but direct studies on the high affinity ferrous iron transporter (Feo transporter) is lacking.

The Feo system plays an essential role in iron acquisition by bacteria in the intestinal tract (16, 23), and mainly consists of FeoA, a small cytoplasmic protein, and FeoB, the membrane-bound permease that is thought to transport ferrous iron into cells; FeoC is also present in some Feo systems. FeoA is thought to interact with the cytoplasmic domain of FeoB to regulate ferrous iron transport (24–27). FeoC is under characterized but was shown to interact with FeoB *in vitro* (25). *C. difficile* encodes three Feo transporter systems of which Feo1 is thought to be the main iron-responsive transporter. Under iron-limiting conditions, *feoB1* is the most transcribed *feo* homolog and is upregulated by >200-fold in mice (11, 28–30). In *C. difficile*, deletion of *feoB1* in non-toxigenic ATCC 700057 did not affect growth in Brain Heart Infusion (BHI) broth, but reduced iron content by approximately 21% and also redox-mediated activity of metronidazole (31). Iron deficiency also perturbs *C. difficile* metabolism, including shifting cells from ferredoxin to flavodoxin-mediated reactions; increases intracellular concentrations of pyruvate and glucose (30, 31); and enhances proline reduction via Stickland metabolism (30). Herein, we show that iron deficiency caused by loss of *feoB1* impacts *C. difficile* toxin production and pathogenesis, and was associated with increased intracellular pyruvate and avirulence in mice.

## RESULTS AND DISCUSSION

### Loss of *feoB1* impairs toxin production

R20291 was adopted as a model strain of epidemic ribotype 027 (32) to delete *feoB1* by allelic exchange to generate R20291Δ*feoB1*; deletion was confirmed by PCR and Sanger sequencing (**Fig S1**). Growth of R20291Δ*feoB1* in BHI broth was not substantially affected, when compared to WT R20291 (**Fig 1A**). Measurement of intracellular iron content, by ICP-OES, revealed a ∼25% reduction in iron content in R20291Δ*feoB1* (0.087 ± 0.006 ppm) compared to the WT (0.116 ± 0.007 ppm) (**Fig 1B**; p ≤0.01, by unpaired t-test). These results are comparable to the *feoB1* deletion mutant of non-toxigenic ATCC 700057, which experienced a ∼21% reduction in intracellular iron when grown in BHI (31). We next quantified toxins in the supernatants of 24 h cultures using an ELISA kit from tgcBIOMICS GmbH. Toxin concentrations in R20291Δ*feoB1*, for TcdA (5.72 ± 3.94 ng/ml) and TcdB (2.50 ± 1.65 ng/ml), were significantly lower than those produced by WT R20291 (227.4 ± 18.24 ng/ml; and 111.2 ± 6.10 ng/ml, respectively) (p<0.0001 and p<0.0001, by one-way ANOVA) (**Fig 1C**). Complementation of R20291Δ*feoB1* with WT *feoB1*, ectopically expressed from its own promoter, increased the synthesis of both toxins (81.04 ± 10.36 ng/ml; and 23.03 ± 5.05 ng/ml, for TcdA and TcdB, respectively (**Fig 1C**) (p<0.0015 and p<0.0041, by one-way ANOVA). Comparable results were attained in cytopathic assays that measure toxin-induced cell rounding of mammalian cells and detects picomolar amounts of toxins (33). Against Vero cells, R20291Δ*feoB1* had toxin titers of ≤10^1^, which was 1000-fold lower than the titers of WT cultures (10^4^ toxin titer; **Fig. 1D**); the complemented strain (R20291Δ*feoB1*_WT *feoB1*) had toxin titers of 10^3^ to 10^4^. These findings were confirmed by automated morphometric analysis of cell rounding to avoid potential bias from manual visualization of cells; as shown in **Fig 1E** 1:1000 diluted supernatants for R20291, R20291Δ*feo*B1 and R20291Δ*feoB1*_WT *feoB1* caused 88.70 ± 1.60, 10.54 ± 0.52 and 52.02 ± 0.74 percentage cell rounding, respectively; noteworthy, cell rounding of Vero cells is more due to TcdB than TcdA (34). Taken together, Δ*feoB1* diminished intracellular iron content and restricted the biosynthesis of TcdA and TcdB, without affecting growth rates in R20291 in BHI.

**Figure 1.**
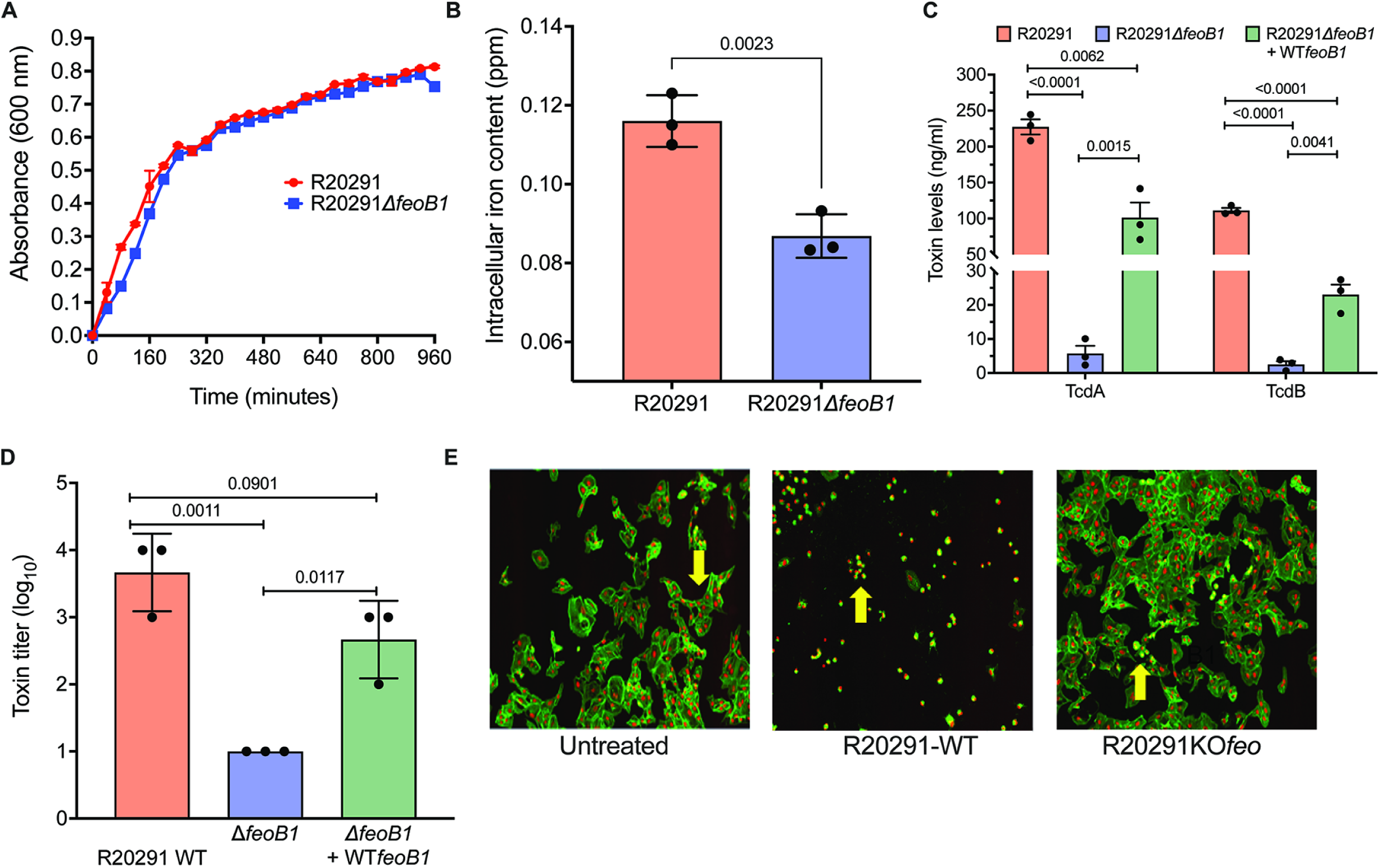
Deletion of *feoB1* attenuates toxin production in *C. difficile* R20291. **A.** Loss of *feoB1* did not affect growth of R20291 in BHI. **B.** Loss of FeoB1 was associated with ∼25% reduction in intracellular iron content in R20291Δ*feoB1*. **C.** TcdA and TcdB toxins were diminished in R20291Δ*feoB1* as quantified by ELISA (limit of detection 0.5 ng/ml for TcdA and TcdB); **D.** Toxin reduction was also confirmed by manual cell rounding analysis, showing decreased toxin titers against Vero epithelial cells (limit of detection 10^1^ dilution); there was ∼ 1000-fold less toxins produced by the mutant. Vero cells are more predictive for cell rounding caused by TcdB. **E.** Cell rounding results were confirmed by automated morphometric analysis. Images show that more cell rounding was more caused by WT R20291 than the mutant at 1:1000 *in situ* dilution. In the untreated control, the yellow arrow shows the morphology of Vero cells, showing actin filaments stained with phalloidin (green) and nuclei stained with DAPI (red); in cells treated with WT supernatants, the yellow arrow shows the morphology of rounded cells. Cells treated with supernatants from the mutant, mostly appeared healthy. Statistical significance in Graphpad Prism was determined by unpaired t-tests with Welch’s correction in b, and by one-way ANOVA with Tukey’s test in c and d.

### *C. difficile* adjust its metabolism to compensate for lack of FeoB1

Ferrous iron is required by enzymes involved in multiple metabolic pathways in *C. difficile*, which that are adopted at various stages of growth (13). Hence, the transcriptome of R20291Δ*feoB1* was analyzed to identify metabolic and virulence-associated pathways that are most affected in mid-logarithmic (OD=0.3), late-logarithmic (OD=0.6) and stationary phase (OD=0.9) stages of growth. Significantly differentially expressed genes (DEGs) were based on p ≤ 0.02 and Log_2_fold change (FC) ≥ 1.5 i.e., FC≥2.8 following analysis in CLC Genomics Workbench version 12 (**Table S1**). **Table 1** highlights selected pathways/genes affected by Δ*feoB1*, including genes that were identified at a less strict criteria of Log_2_fold change (FC) ≥ 0.58 i.e., FC≥1.5. Across the three growth phases, there were 435 common differentially expressed genes, of which 151 were upregulated and 284 were downregulated (**Fig S1**). The transcriptome data revealed Δ*feoB1* affected redox and flavin-dependent processes (**Table 1**; **Fig. 2A-C**; **Fig. S1**). Although our studies were done in BHI media, expression changes seen in our RNAseq were generally comparable to those reported for a *C. difficile* CD630 *fur* mutant that was grown in low iron media (30), including carbohydrate metabolism, iron homeostasis, pathogenesis, flagellar proteins, cell wall biosynthesis genes (**Table 1**). As described below, we focused on gene signatures that are linked to iron metabolism and *C. difficile* pathogenesis, indicating how *C. difficile* coped with reduced intracellular iron content.

**Figure 2.**
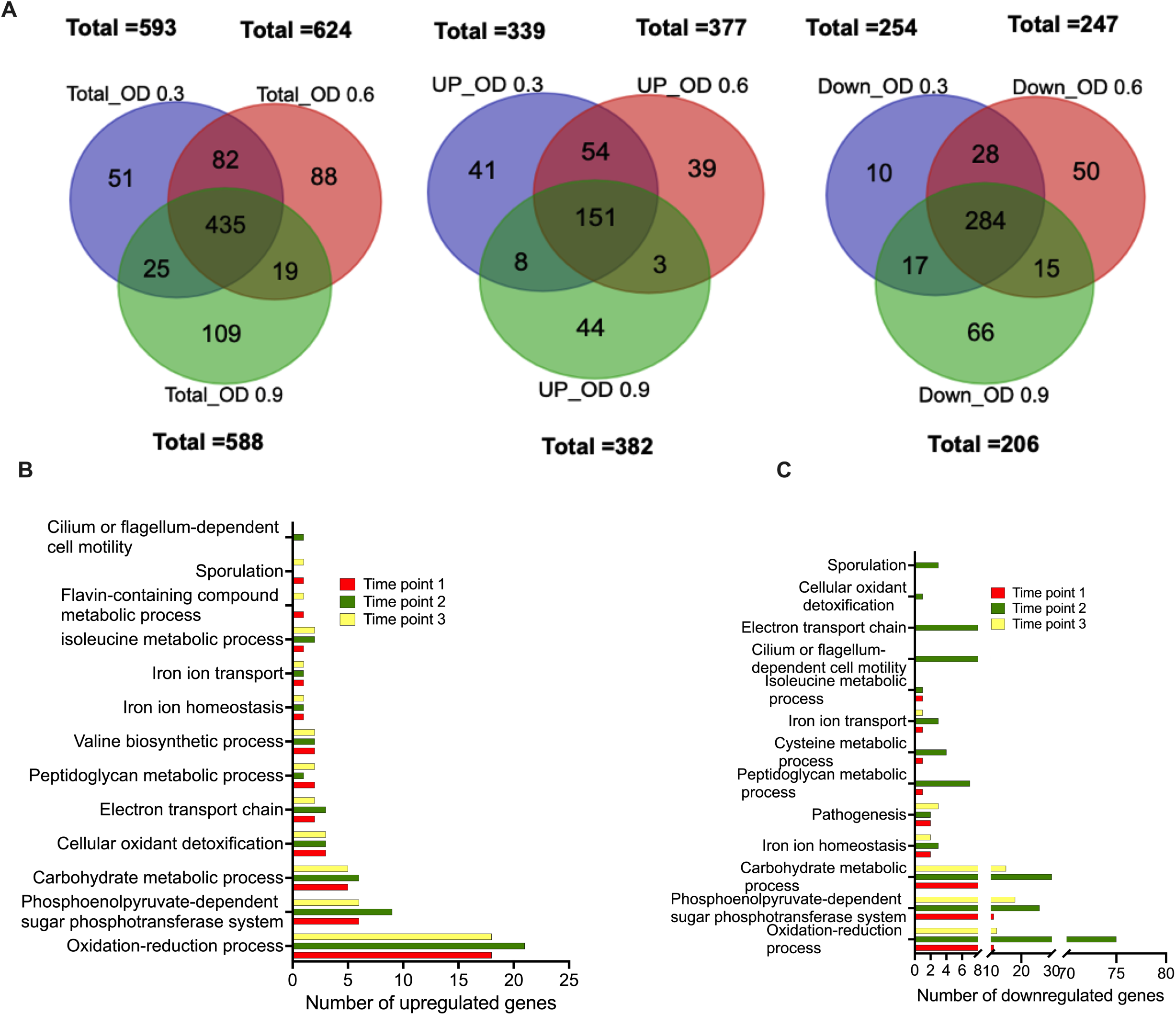
Transcriptome changes associated with lack of *feoB1* in R20291. **A.** Venn diagrams show total differentially expressed genes (DEGs) and those that were upregulated or downregulated at different growth phases (mid logarithmic OD600nm=0.3; late-logarithmic OD600nm=0.6; and stationary phase OD600nm=0.9). **B, C**. Gene Ontology analysis of DEGs identified different metabolic processes altered by loss of *feoB1*, including iron homeostasis, pathogenesis, and redox reactions.

**Table 1.**
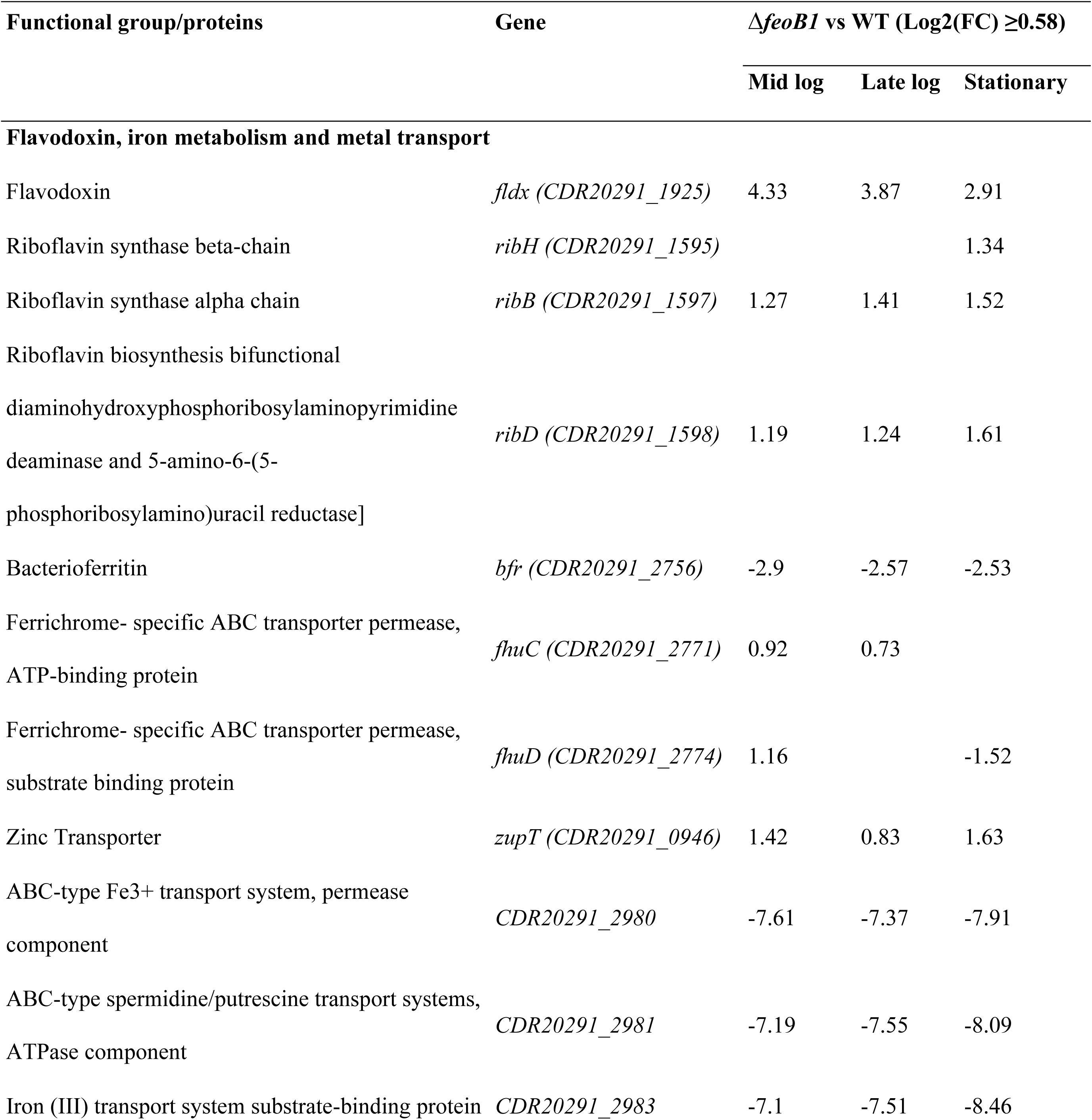
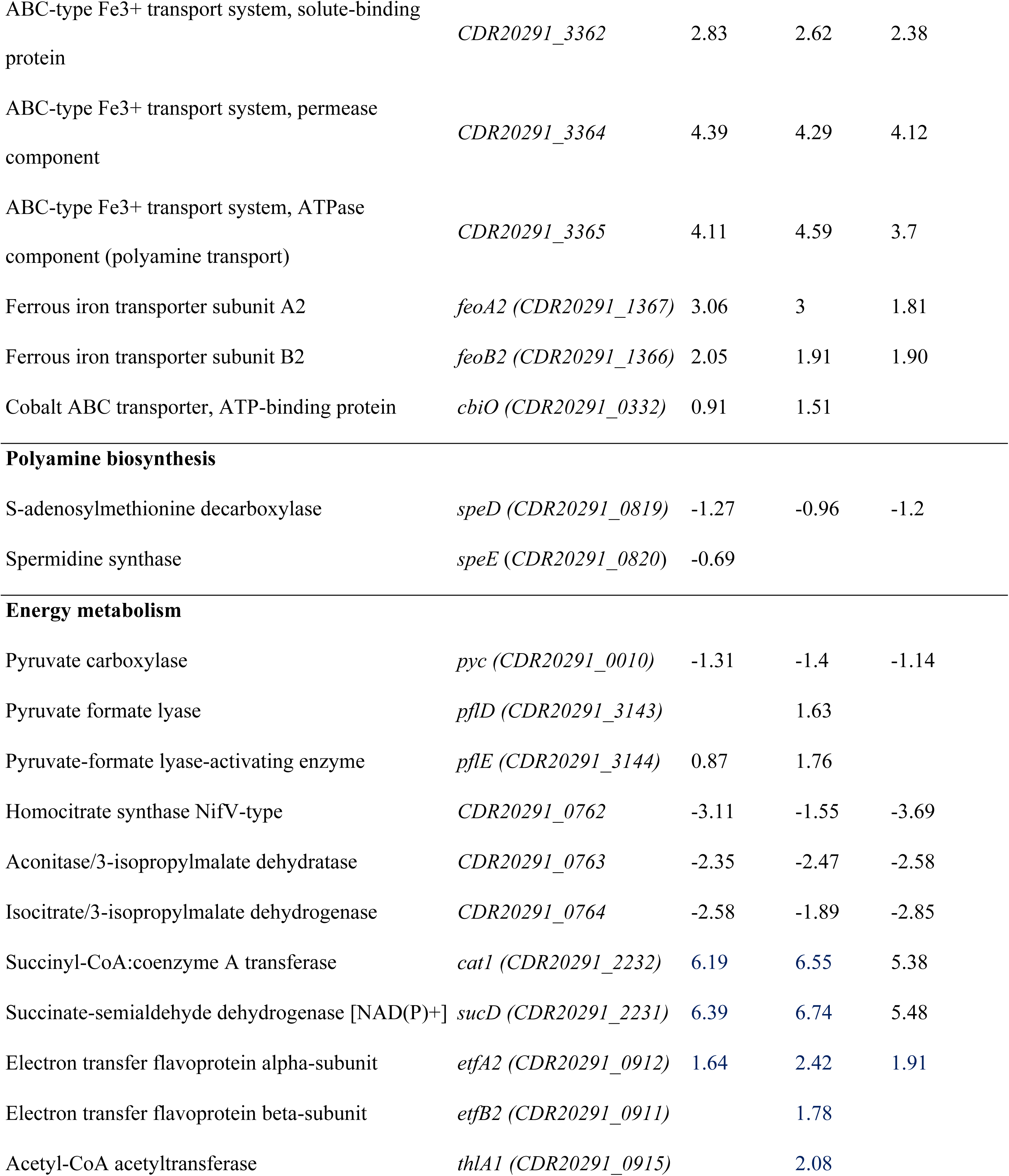
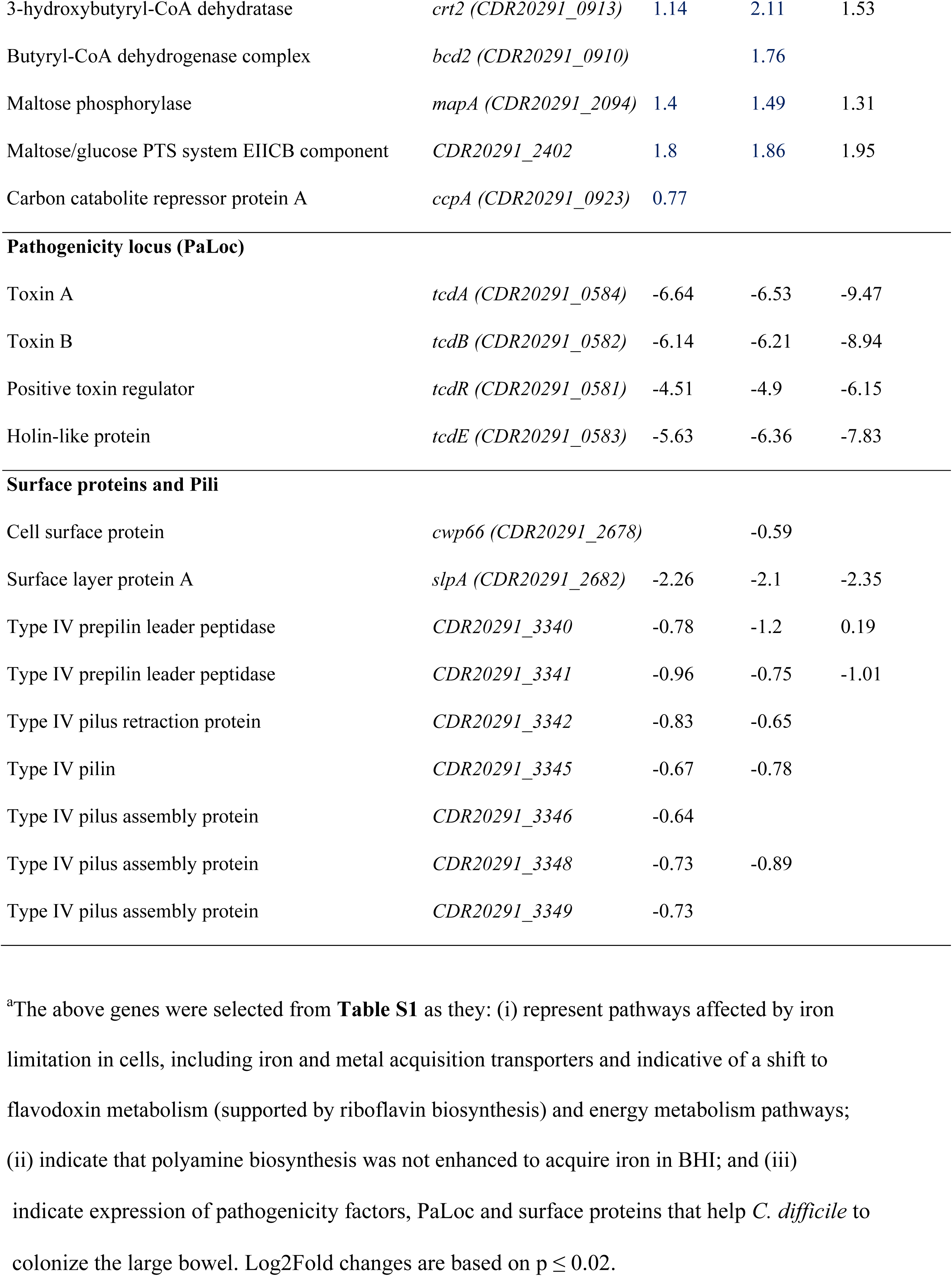
Selected genes^a^ in *C. difficile* R20291 that were differentially expressed in R20291Δ*feoB1* compared to WT R20291.

| Functional group/proteins | Gene | $\Delta feoB1$ vs WT (Log2(FC) $\geq 0.58$) | | |
| --- | --- | --- | --- | --- |
|  |  | Mid log | Late log | Stationary |
| Flavodoxin, iron metabolism and metal transport |  |  |  |  |
| Flavodoxin | *fldx* (CDR20291_1925) | 4.33 | 3.87 | 2.91 |
| Riboflavin synthase beta-chain | *ribH* (CDR20291_1595) |  |  | 1.34 |
| Riboflavin synthase alpha chain | *ribB* (CDR20291_1597) | 1.27 | 1.41 | 1.52 |
| Riboflavin biosynthesis bifunctional<br>diaminohydroxyphosphoribosylaminopyrimidine<br>deaminase and 5-amino-6-(5-phosphoribosylamino)uracil reductase] | *ribD* (CDR20291_1598) | 1.19 | 1.24 | 1.61 |
| Bacterioferritin | *bfr* (CDR20291_2756) | -2.9 | -2.57 | -2.53 |
| Ferrichrome- specific ABC transporter permease,<br>ATP-binding protein | *fhuC* (CDR20291_2771) | 0.92 | 0.73 |  |
| Ferrichrome- specific ABC transporter permease,<br>substrate binding protein | *fhuD* (CDR20291_2774) | 1.16 |  | -1.52 |
| Zinc Transporter | *zupT* (CDR20291_0946) | 1.42 | 0.83 | 1.63 |
| ABC-type Fe3+ transport system, permease<br>component | *CDR20291_2980* | -7.61 | -7.37 | -7.91 |
| ABC-type spermidine/putrescine transport systems,<br>ATPase component | *CDR20291_2981* | -7.19 | -7.55 | -8.09 |
| Iron (III) transport system substrate-binding protein | *CDR20291_2983* | -7.1 | -7.51 | -8.46 |
| ABC-type Fe<sup>3+</sup> transport system, solute-binding protein | *CDR20291_3362* | 2.83 | 2.62 | 2.38 |
| ABC-type Fe<sup>3+</sup> transport system, permease component | *CDR20291_3364* | 4.39 | 4.29 | 4.12 |
| ABC-type Fe<sup>3+</sup> transport system, ATPase component (polyamine transport) | *CDR20291_3365* | 4.11 | 4.59 | 3.7 |
| Ferrous iron transporter subunit A2 | *feoA2 (CDR20291_1367)* | 3.06 | 3 | 1.81 |
| Ferrous iron transporter subunit B2 | *feoB2 (CDR20291_1366)* | 2.05 | 1.91 | 1.90 |
| Cobalt ABC transporter, ATP-binding protein | *cbiO (CDR20291_0332)* | 0.91 | 1.51 |  |
| Polyamine biosynthesis |  |  |  |  |
| S-adenosylmethionine decarboxylase | *speD (CDR20291_0819)* | -1.27 | -0.96 | -1.2 |
| Spermidine synthase | *speE (CDR20291_0820)* | -0.69 |  |  |
| Energy metabolism |  |  |  |  |
| Pyruvate carboxylase | *pyc (CDR20291_0010)* | -1.31 | -1.4 | -1.14 |
| Pyruvate formate lyase | *pflD (CDR20291_3143)* |  | 1.63 |  |
| Pyruvate-formate lyase-activating enzyme | *pflE (CDR20291_3144)* | 0.87 | 1.76 |  |
| Homocitrate synthase NifV-type | *CDR20291_0762* | -3.11 | -1.55 | -3.69 |
| Aconitase/3-isopropylmalate dehydratase | *CDR20291_0763* | -2.35 | -2.47 | -2.58 |
| Isocitrate/3-isopropylmalate dehydrogenase | *CDR20291_0764* | -2.58 | -1.89 | -2.85 |
| Succinyl-CoA:coenzyme A transferase | *cat1 (CDR20291_2232)* | 6.19 | 6.55 | 5.38 |
| Succinate-semialdehyde dehydrogenase [NAD(P)+] | *sucD (CDR20291_2231)* | 6.39 | 6.74 | 5.48 |
| Electron transfer flavoprotein alpha-subunit | *etfA2 (CDR20291_0912)* | 1.64 | 2.42 | 1.91 |
| Electron transfer flavoprotein beta-subunit | *etfB2 (CDR20291_0911)* |  | 1.78 |  |
| Acetyl-CoA acetyltransferase | *thlA1 (CDR20291_0915)* |  | 2.08 |  |
| 3-hydroxybutyryl-CoA dehydratase | *crt2 (CDR20291_0913)* | 1.14 | 2.11 | 1.53 |
| Butyryl-CoA dehydrogenase complex | *bcd2 (CDR20291_0910)* |  | 1.76 |  |
| Maltose phosphorylase | *mapA (CDR20291_2094)* | 1.4 | 1.49 | 1.31 |
| Maltose/glucose PTS system EIICB component | *CDR20291_2402* | 1.8 | 1.86 | 1.95 |
| Carbon catabolite repressor protein A | *ccpA (CDR20291_0923)* | 0.77 |  |  |
| Pathogenicity locus (PaLoc) |  |  |  |  |
| Toxin A | *tcdA (CDR20291_0584)* | -6.64 | -6.53 | -9.47 |
| Toxin B | *tcdB (CDR20291_0582)* | -6.14 | -6.21 | -8.94 |
| Positive toxin regulator | *tcdR (CDR20291_0581)* | -4.51 | -4.9 | -6.15 |
| Holin-like protein | *tcdE (CDR20291_0583)* | -5.63 | -6.36 | -7.83 |
| Surface proteins and Pili |  |  |  |  |
| Cell surface protein | *cwp66 (CDR20291_2678)* |  | -0.59 |  |
| Surface layer protein A | *slpA (CDR20291_2682)* | -2.26 | -2.1 | -2.35 |
| Type IV prepilin leader peptidase | *CDR20291_3340* | -0.78 | -1.2 | 0.19 |
| Type IV prepilin leader peptidase | *CDR20291_3341* | -0.96 | -0.75 | -1.01 |
| Type IV pilus retraction protein | *CDR20291_3342* | -0.83 | -0.65 |  |
| Type IV pilin | *CDR20291_3345* | -0.67 | -0.78 |  |
| Type IV pilus assembly protein | *CDR20291_3346* | -0.64 |  |  |
| Type IV pilus assembly protein | *CDR20291_3348* | -0.73 | -0.89 |  |
| Type IV pilus assembly protein | *CDR20291_3349* | -0.73 |  |  |

<sup>a</sup>The above genes were selected from **Table S1** as they: (i) represent pathways affected by iron limitation in cells, including iron and metal acquisition transporters and indicative of a shift to flavodoxin metabolism (supported by riboflavin biosynthesis) and energy metabolism pathways; (ii) indicate that polyamine biosynthesis was not enhanced to acquire iron in BHI; and (iii)
indicate expression of pathogenicity factors, PaLoc and surface proteins that help *C. difficile* to colonize the large bowel. Log2Fold changes are based on $p \leq 0.02$ .

### Iron-responsive gene transcription

Iron starvation in *C. difficile* increases the transcription of flavodoxin (*fldx*), ferric uptake regulator (*fur*), riboflavin biosynthesis and alternate iron uptake transporters (30). Throughout the growth of R20291Δ*feoB1* there was upregulation of the flavodoxin encoded by *CDR20291_1925*; riboflavin biosynthesis genes *ribB* and *ribD*; and downregulation of *CDR20291_2756*, encoding the iron storage protein bacterioferritin (**Table 1**; **Fig. 3A**). Upregulation of flavodoxin and riboflavin biosynthesis genes reflect a metabolic shift to adopt flavin cofactors, to likely substitute for the iron-dependent ferredoxin (35, 36). Downregulation of bacterioferritin suggests cells sought to increase cellular availability of iron, by decreasing storage of ferric iron in bacterioferritin. Interestingly, the expressions of *fezA* and *fezB* (*CDR20291_1517* and *CDR20291_1516*) were not substantially affected throughout growth, which contrasts with observations in CD630, where iron limiting conditions caused by bipyridyl exposure reduced mid-logarithmic growth (37). However, in our study, growth in BHI was not attenuated, nor was iron limited, allowing us to map effects that are not due to significant growth defects. However, alternate iron acquisition transporters were also upregulated such as ferrichrome-specific ABC transporter genes (*fhuC* and *fhuD*) (**Fig. 3A**), the low-affinity zinc transporter encoded by *zupT* and Feo2 encoded by *feoA2* and *feoB2* genes (**Table 1**). Increased expression of *zupT* also occurred in *C. difficile* CD630 under bipyridyl-induced iron limiting conditions (37) but it is unknown whether ZupT transports iron as previously shown in *Escherichia coli* (20). Deletion of *zupT* in *C. difficile* was reported to impair colonization in mice, although its effect on toxin production was not described (38).

**Figure 3.**
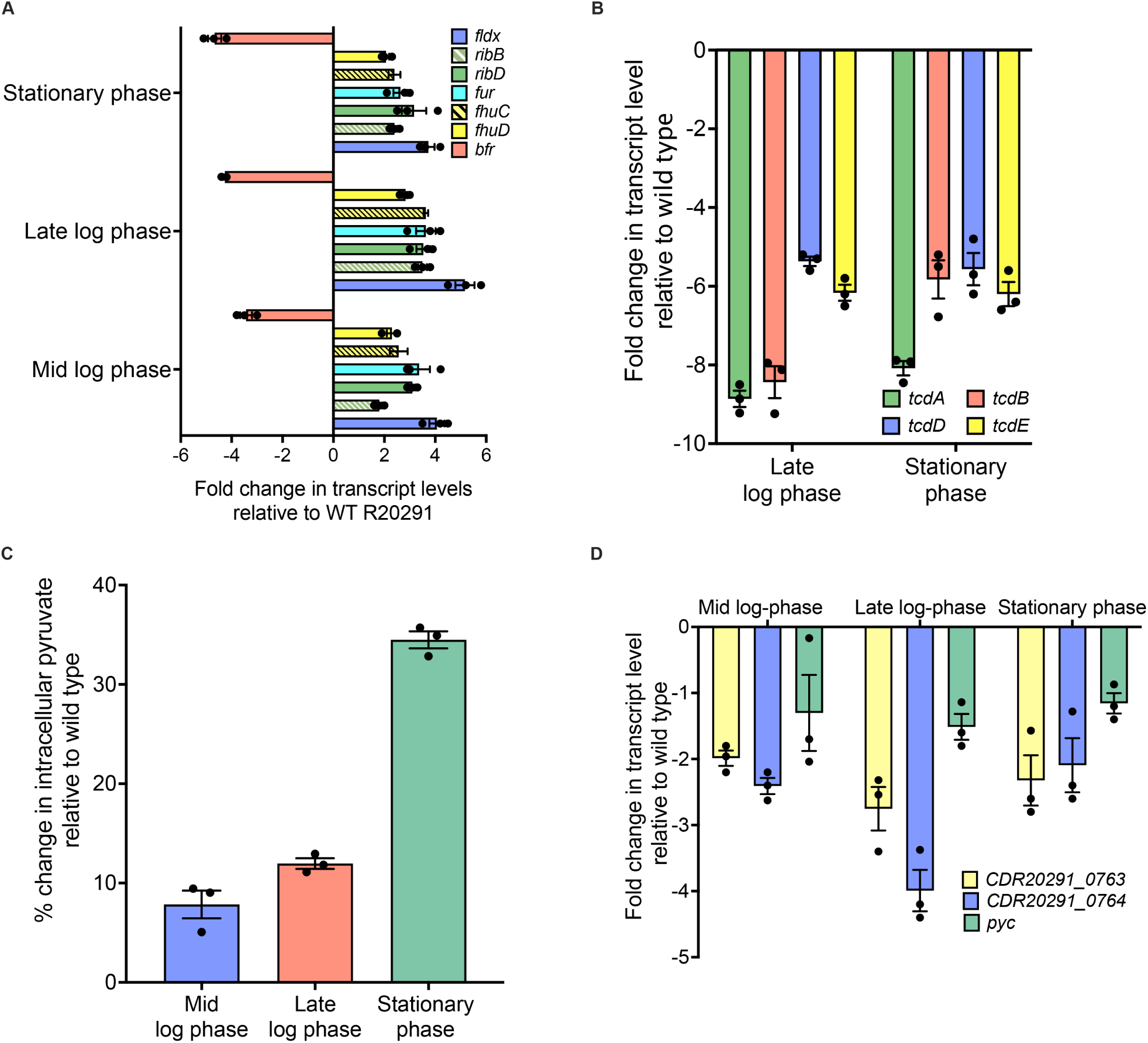
Effect of loss of FeoB1 on transcription of iron-responsive genes, PaLoc, TCA and pyruvate accumulation. **A.** Across three growth phases, loss of *feoB1* dysregulated iron-responsive genes in R20291Δ*feoB1*, shown by upregulation of flavodoxin (*fldx*), ferrichrome-specific ABC transporter permease (*fhuC* and *fhuD*), ferric uptake regulator (*fur*) and riboflavin biosynthesis genes (*ribB* and *ribD*), and downregulation of bacterioferritin (*bfr*). **B.** Downregulation of toxins genes (*tcdA*, *tcdB*, *tcdD* and *tcdE*) was seen in R20291Δ*feoB1* late-logarithmic (OD600nm=0.6) and stationary phase cultures (OD600nm=0.9). **C.** Across the three growth phases intracellular pyruvate was higher in R20291Δ*feoB1* (1.25 ± 0.14 nm, 1.55 ± 0.11 nm and 2.24 ± 0.16 nm, respectively) compared to R20291 (1.16 ± 0.13 nm, 1.39 ± 0.08 nm and 1.66 ± 0.11 nm, respectively). **D.** Downregulation of genes encoding enzymes of the oxidative branch of the Krebs cycle was also seen (*pyc* encodes pyruvate carboxylase; *CDR20291_0763* aconitase; and *CDR20291_0764* encodes isocitrate/3-isopropylmalate dehydrogenase).

### Relationship between FeoB1, pyruvate metabolism and toxin production

In R20291Δ*feoB1* (**Fig 3B**) PaLoc genes were significantly downregulated, including *tcdA* and *tcdB*, the sigma factor *tcdR* and holin *tcdE* (**Table 1**). Thus, the low levels of TcdA and TcdB in supernatants from R20291Δ*feoB1* directly correlate with reduced expression of *tcdA* and *tcdB*, suggesting loss of *feoB1* enhanced their regulatory control. We recently reported that Δ*feoB1* and disruption of *iscR* encoding iron-sulfur cluster regulator caused pyruvate accumulation in non-toxigenic ATCC 700057 (31). Since pyruvate build-up negatively impacts toxin production (11), we measured this metabolite in R20291Δ*feoB1* and found that it accumulated throughout all three growth phases, when compared to R20291 (**Fig 3C**). This was especially pronounced in the stationary phase, having a 34.48% increase in pyruvate, compared to 7.84% and 11.94% in mid-and late-logarithmic cultures, respectively. Pyruvate feeds into multiple metabolic pathways, where it can be converted to formate by pyruvate-formate lyase (Pfl); lactate by lactate dehydrogenase (Ldh); acetyl-CoA by pyruvate-ferredoxin oxidoreductase (NifJ); or enter the incomplete Krebs cycle of *C. difficile* to be converted to oxaloacetate by pyruvate carboxylase (Pyc) (13). Many of these genes were differentially regulated in the transcriptome (**Table 1**). In cultures at OD ∼0.6 *pflD* and *pflE* were increased by 3.09-and 3.39-fold, respectively but were not significantly differentially expressed in stationary phase cultures. *C. difficile* encodes an incomplete Krebs cycle, in which pyruvate enters the oxidative branch via Pyc, which converts it to oxaloacetate. Oxaloacetate is then metabolized to citrate, likely by homocitrate synthase (NifV, CDR20291_0762), followed by conversion to isocitrate and 2-oxoglutarate, catalyzed by aconitase (CDR20291_0763) and isocitrate/3-isopropylmalate dehydrogenase (CDR20291_0764), respectively (39). Interestingly, *pyc* was approximately two-fold downregulated (-2.2 to-2.64) throughout the growth cycle. *CDR20291_0762* to *CDR20291_0764* were also all downregulated in the transcriptome (-1.89 to 12.93-fold across the three growth phases (**Table 1**). RT-qPCR of *pyc*, *CDR20291_0763* and *CDR20291_0764* confirmed their downregulation (**Fig. 3D**). Thus, it is likely that downregulation of *pyc* and the oxidative branch of the Kreb cycle contributed to the build-up of pyruvate.

### Deletion of *feoB1* attenuated virulence in mice with colitis CDI

Given that loss of *feoB1* elevated pyruvate levels and attenuated toxin production *in vitro*, we examined the virulence of R20291Δ*feoB1* in a model of colitis CDI in C57BL/6 showing that loss of *feoB1* attenuated pathogenesis (**Fig. 4A-D**). Since this model is characterized by intestinal bleeding it enabled assessment of R20291Δ*feoB1* virulence in a setting that favors iron availability in the large bowel. Infection with WT R20291 resulted in half of the mice becoming moribund, whereas all fifteen mice infected with R20291Δ*feoB1* survived (**Fig. 4C**); p=0.0003 according to log rank test. Significant weight loss was also observed in animals infected with R20291 (**Fig. 4D**), whereas mice infected with R20291Δ*feoB1* either gained weight or experienced marginal weight loss (p=0.0009 by unpaired t-test). At sacrifice, mice infected with R20291 exhibited inflamed ceca, while those infected with R20291Δ*feoB1* had ceca resembling the uninfected mice (**Fig. 4B**). Consistent with these observations, colonization rates were significantly higher in mice infected with R20291, indicating poor infectivity by R20291Δ*feoB1*. In moribund mice, spore bioburdens were ∼10^7^ CFU/g for R20291, which was ∼2 logs [∼99%] greater than the bioburdens from animals that survived infection with R20291 (p<0.0001 by unpaired t-test; **Fig. 4E**). However, animals infected with R20291Δ*feoB1* had even lower bioburdens ∼10^4^ CFU/g (p<0.0001 by ANOVA; **Fig. 4E**). As a control, there was no difference in *in vitro* germination between R20291 and R20291Δ*feoB1* (**Fig. S2**). Furthermore, cecal isolates from the respective animals were confirmed to be R20291 from PCR amplicons of *CDR20291_1801* on *Tn6192* transposon in R20291 and *tcdA*; R20291Δ*feoB1* was further confirmed from the absence of *feoB1* (**Fig. S3**).

**Figure 4.**
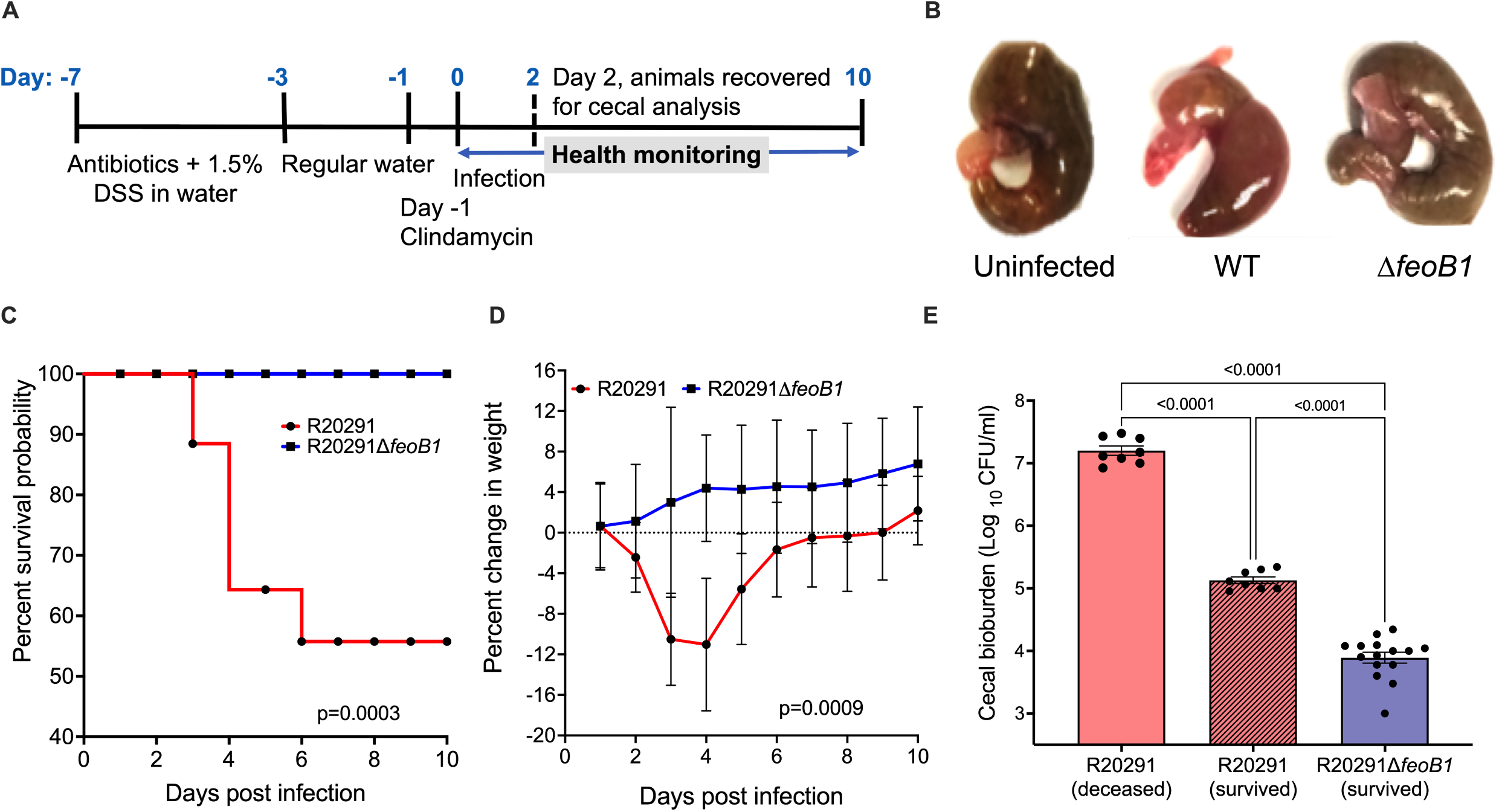
Deletion of FeoB1 attenuates colonization and virulence in mice. **A.** Experimental summary of the colitis CDI model used to evaluate the virulence of R20291 and R20291Δ*feoB1*. **B.** Ceca from the mice infected with the wild type were red and inflamed in contrast to ceca from the *feoB1* mutant that resembled the uninfected control. **C.** Kaplan-Meier survival analysis show significant difference (p=0.0003) between mortality risks of mice infected with R20291 compared to R20291Δ*feoB1*. **D.** Animals infected with R20291 had significant weight loss (p=0.0009) compared to R20291Δ*feoB1* infected mice. **E**. Cecal bioburdens were highest in deceased mice infected with R20291 than mice surviving infection, while R20291Δ*feoB1* infected mice had lowest bioburdens. Data are based on group numbers of n=16 mice for R20291 and n=15 mice for R20291Δ*feoB1* (a mouse from the R20291Δ*feoB1* group was removed due to inadvertent infection with R20291). All statistical analyses were done in GraphPad prism 10; including Log-rank statistical testing in c; unpaired t-tests with Welch’s correction in d; and one-way ANOVA with alpha 0.05 in e.

### Comparison of pathogenesis in mice at two days of infection

Across different mouse models of CDI, infection on day 2 produces significant morbidity allowing capture of the acute response to infection. We therefore conducted a separate study in which mice were culled at 48 h to examine effects on animal health, immune response and spore and toxin burdens. Mice infected with the WT had significant weight loss compared to the pre-infected weights (**Fig. 5A**; p ≤0.01), whereas those infected with the mutant gained weight (p ≤0.05). For R20291Δ*feoB1* cecal bioburdens were almost 3-logs less than mice infected with R20291, which was also evident in fecal bioburdens at 24 and 48 h of post-infection (**Fig. 5B, C**); unpaired t-test p<0.0001). Cecal toxins were also reduced in mice infected with R20291Δ*feoB1* and were 1-2 logs (90-99%) less than mice infected with the WT infected, which had toxin titers of 10^2^-10^3^ per gram of cecal content (**Fig. 5D**). Measurement of inflammatory responses, by qRT-PCR on cecal tissue, showed significantly increased expression of pro-inflammatory tumor necrosis factor-alpha (TNF-α), granulocyte-macrophage colony-stimulating factor (GM-CSF) and anti-inflammatory interleukin-10 (IL-10) in mice infected with the WT compared to R20291Δ*feoB1* (**Fig. 5E**). While TNF-α and GM-CSF promote neutrophil recruitment in CDI, the upregulation of IL-10 may occur to repress TNF-α and GM-CSF (40). Elevation of these three cytokines may be markers for CDI severity in the CDI colitis model (41). Overall, these findings indicate that *C. difficile* requires FeoB1 to colonize and produce toxin-mediated disease symptoms.

**Figure 5.**
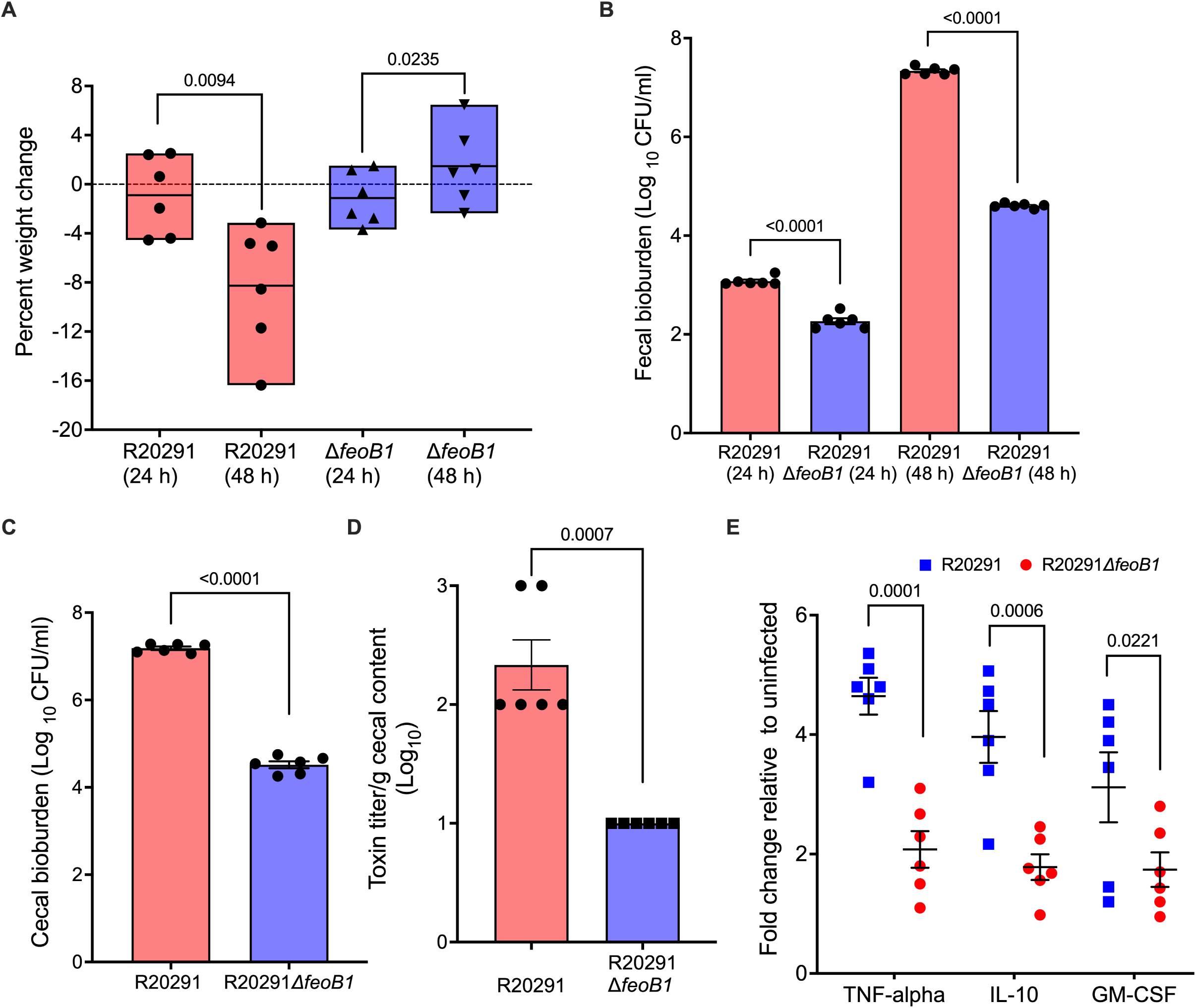
Comparative analysis of virulence in mice at 48 h post-infection. **A.** Significant weight loss was seen 48 hours after infection with R20291 whereas mice infected with R20291Δ*feoB1* gained weight, which was also significant. **B, C.** Reduced colonization by R20291Δ*feoB1* was also evident from bioburdens analyzed from the ceca taken at 48 hours and fecal samples recovered at 24 and 48 h. **D, E.** Lower toxin titers and expression of cytokine in cecal tissues were also evident. Statistical significance was determined in GraphPad Prism 10.0 by unpaired t-tests with Welch’s correction in a-d and in e by two-way ANOVA with Šídák’s multiple comparisons test.

## CONCLUSION

Our study clearly shows that *feoB1* is necessary for *C. difficile* infectivity and virulence in mice, which is a finding that mirrors the loss of ferrosome storage upon deletion of *fezA* (22). Noteworthy, we adopted BHI and the colitis CDI model to evaluate the responses of *C. difficile* in environments that are available iron sources. Although FeoB1 may be the main permease for ferrous iron acquisition, its deletion did not affect *in vitro* growth, as *C. difficile* likely had different iron sources in BHI media and could remodel its metabolism to utilize flavodoxin cofactors and less iron-dependent energy generation involving coupling of the ATP synthetase and Rnf complex (30). Upregulation of the proline reductase operon in the transcriptome of mid-log cultures (**Table S1**) might also indicate that loss of *feoB1* increased the need for the Stickland metabolism, in which the oxidation of an amino acid (e.g., cysteine) is coupled to the reduction of another (e.g., proline) to produce ATP (13, 14).

The Feo system is also essential for other intestinal pathogens such as *E. coli*, *Helicobacter pylori* (42) and *Salmonella typhimurium* (23), which encode a single FeoB homolog. For example, without the Feo system *S. typhimurium* is unable to successfully colonize the large intestine of mice, but it can grow in the circulatory system and systemic body sites (23). Similarly, *H. pylori* requires FeoB to colonize the gastric mucosa of mice, despite encoding transporters for ferric iron and an ability to acquire iron from lactoferrin and transferrin (42, 43). This reports, along with findings of this study, indicate that FeoB1 is required for pathogenesis in acid and anaerobic environments along the gastrointestinal tract. Hence, as the large bowel is the only site for CDI, our findings indicate FeoB1 is indispensable for CDI disease, even if other forms of iron are available in the colon during infection or *C. difficile* expresses other ferrous iron transporters (e.g., FeoB2, FeoB3 or ZupT). This contrasts with ferric hydroxamate transporter that is dispensable for colonization and virulence in mice (21). Nonetheless, research is needed to establish the cellular functions of *C. difficile* FeoB2 and FeoB3, and the structure-function relationships of all three Feo systems in *C. difficile*. Beyond impacting toxin production, iron starvation also hinders the expression of other colonization determinants in *C. difficile*. For example, iron starvation downregulated pili and flagella associated genes of a CD630 *fur* mutant (30). Our transcriptome indicated Type IV pilin, surface layer protein A (*slpA*) and adhesin encoded by *cwp66* were downregulated (**Table 1**). In *C. difficile*, Type IV pilin contributes to adherence to epithelial cells and cell-cell attachment in biofilms; it was reported that deletion of the Type IV pilin gene *pilB1* (*CDR20291_3349*), resulted in poorer colonization of mice (44). Similarly, loss of *slpA* also attenuates *C. difficile* pathogenesis in animal models (45). Hence, loss of *feoB1* imposes different pleiotropic effects on pathogenesis that may involve reduced uptake of ferrous iron, poor toxin production and poor colonization of the gut epithelia. Taken together, these results support the potential for FeoB1 to become a therapeutic target for the discovery of agents that alter *C. difficile* metabolism to block the biosynthesis of TcdA and TcdB and surface proteins required for colonization.

## MATERIALS AND METHODS

### Strains and culture conditions

*C. difficile* R20291, R20291Δ*feoB1* and the complemented strain were routinely grown in pre-reduced BHI broth or agar at 37°C in a Whitley A35 anaerobic workstation (Don Whitley Scientific). *E. coli* SD46 was grown at 37°C in Luria– Bertani (LB) broth or agar. D-cycloserine (250 μg/ml), cefoxitin (8 μg/ml), and thiamphenicol (15 μg/ml) were used to selectively culture *C. difficile* containing plasmids, whereas chloramphenicol (15 μg/ml) and ampicillin (50 μg/ml) were used to grow *E. coli* SD46. For cell rounding assays, Vero epithelial cells were cultured in 10% fetal bovine serum and Dulbecco’s Modified Eagle Medium at 37°C with 5% carbon dioxide in a humidified incubator.

### Growth kinetics

Overnight cultures were diluted 1:100 and grown to mid-logarithmic phase before being added to 96 - well plates containing fresh pre-reduced BHI. OD600 values were automatically recorded under anaerobic conditions in a Synergy H1 microplate reader (BioTek) over 16 h.

### Genetic manipulation of *C. difficile*

R20291Δ*feoB1* was generated by allelic exchange, exactly as we described (19). Essentially, *E. coli* SD46 carrying the *feoB1* allelic deletion cassette in pMTL-SC7215 was conjugated into R20291, before subjecting the recombinant strain to serial passage to select for double crossover mutants. The deletion was confirmed by PCR and Sanger sequencing. R20291Δ*feoB1* was complemented with WT *feoB1* expressed from its promoter in pMTL84151, as we described (19). The strain was maintained on pre-reduced BHI supplemented with thiamphenicol (15 μg/ml). The primers used in the study are listed in **Table S2**.

### Quantification of *C. difficile* toxins

Strains were grown anaerobically in pre-reduced BHI at 37°C for 24 h. Samples were centrifuged at 5000 rpm for 5 min and supernatants recovered for ELISA and cell rounding assays, as follows. **(i) ELISA quantification.** 100 µl of diluted sample was used to measure TcdA and TcdB with the tgcBIOMICS Gmbh ELISA kit that quantifies both toxins, against standard curves from purified TcdA and TcdB, according to the manufacturer’s instructions. **(ii) Standard cell rounding assay.** Toxin titers were determined against Vero epithelial cells (∼10^5^ cells/ml) that were adhered overnight. Supernatants from neat and various diluted samples were added to the cells and incubated for 4 h. Cytopathic effects (cell rounding) were visualized by microscopy, using six views per well. Toxin titers were defined as the lowest dilution in which cell rounding was no longer visualized. (**iii) Determination of cell rounding by morphometric analysis.** Briefly, 50 µl of 500-1000 Vero epithelial cells (ATCC) were seeded per well of a 384-well black plate and incubated for ∼18 h. From a source plate containing diluted TcdB to generate a standard curve and undiluted culture supernatants 0.05 µl of samples were automatedly transferred to the 384-well plate containing the Vero cells using an Echo acoustic liquid handler (Labcyte). This represented an *in situ* dilution of 1:1000 of the culture supernatants. Samples were incubated at 37°C for 6 h in 5% carbon dioxide, followed by addition of 10 µl of 25% (v/v) paraformaldehyde. Cells were fixed overnight at 4°C, washed with PBS and stained with the nucleic acid stain DAPI (25 µl). After incubation in the dark for 15 min at room temperature, they were stained with Alexa Fluor 488 phalloidin strain (40 µl) at a concentration of 0.0006 μM in PBS with 5% milk powder solution. After staining in the dark for 2 h, cells were washed twice, and PBS (50 µl) was added. Cell images were captured using IN Cell Analyzer 6000 from GE Healthcare, with DAPI (excitation 358 nm and emission 451 nm) and phalloidin (excitation 495 nm and emission 518 nm). Images were then processed as described previously (8), to capture morphometric features and classify cells into control-like or toxin-treated-like (small, rounded) phenotypes to establish the proportion of toxin-treated-like cells per well.

### Quantification of intracellular iron by ICP-OES

Overnight grown cells were subcultured (1:100) in fresh BHI broth and harvested after 6 h for ICP-OES analysis to determine the cellular iron levels as we previously described (31).

### Quantification of intracellular pyruvate

This assay was performed as we previously described (31); essentially harvesting samples at OD∼0.3, 0.6 and 0.9 to measuring intracellular pyruvate in cell lysates using the Pyruvate Colorimetric/Fluorometric Assay kit from Biovision.

### RNA Sequencing and analysis

Triplicates of R20291 and R20291Δ*feoB1* were grown overnight in pre-reduced BHI broth. The next day, cultures were diluted 1:100 in pre-reduced BHI broth and grown in the anaerobic chamber. Approximately 20 ml samples of each strain were harvested at three-time points OD∼0.3, ∼0.6, and ∼0.9 and further aliquots of each time point also stored for subsequent analyses. These time points were selected to assess changes in gene transcription at different points in the bacterial growth cycle. Bacterial RNAprotect™ reagent (QIAGEN) was added and RNA isolated using Qiagen’s RNeasy mini kit. RNA quality assessment, library preparation and Hi-Seq 2 x150 paired-end sequencing were performed by GENEWIZ (New Jersey, USA).

### Analysis of RNA sequencing data

CLC Genomics Workbench version 12.0 was used to analyze RNA sequencing data. After trimming the adapters, the data was checked for its quality and relatedness using Principal Component Analysis (PCA). Further, *C. difficile* R20291 coding sequence (CDS, GenBank accession number: FN545816.1) was used for alignment and analysis. Differential gene expression of R20291 Δ*feoB1* was analyzed against R20291 wild type. Functional clustering of DEGs was done using KEGG Mapper. The raw RNA-Seq data are deposited in NCBI database under accession number PRJNA753147.

### Transcriptional analysis by RT-qPCR

RNA isolated from *C. difficile* samples was converted to cDNA by M-MLV reverse transcriptase (Quantabio) using qScript cDNA SuperMix. qScript One-Step SYBR Green qRT-PCR Kit, ROX (Quantabio) and gene-specific primers were used to amplify genes in Applied Biosystems ViiA7 real-time PCR machine. Transcript levels were calculated by the comparative Ct method (ΔΔCT method) and data normalized to 16S rRNA (46). For analysis of cytokine levels in the cecal tissues, the same Qiagen’s RNeasy mini kit was used to isolate RNA and qRT-PCR was carried out as above. The data was normalized to the beta-actin gene.

### Mouse model of colitis CDI

These experiments were conducted under an animal use protocol approved by the Institutional Animal Care and Use Committee at Texas A&M University. The model of CDI was modified from that described by Zhou et al. (41). Male and female C57/BL6 mice (6 weeks) were from Envigo. For 5 days, mice were given water containing kanamycin (0.4 µg/ml), gentamicin (0.035 µg/ml), colistin (850 U/ml), metronidazole (0.215 µg/ml), vancomycin (0.045 µg/ml) and 1.5% w/v of dextran sodium sulfate. This was followed by two days of regular drug-free water. On day-1, mice were injected (i.p.) with clindamycin (10 mg/kg). Approximately 20 h later (day 0) they were infected with 10^5^ spores of either R20291 or R20291Δ*feoB1*. Three sets of experiments were carried out, as summarized in the results. *C. difficile* spores were prepared according to a standard protocol (47). After infection, fecal samples were collected daily, and ceca recovered from euthanized moribund animals or at the experimental endpoint. Animals were monitored three times per day and scored for: health appearance (hunched posture, ruffled fur, and diarrhea, and wet tail); weight loss; and decreases in body temperature (**Table S3**).

### Analysis of cecal and fecal bioburdens

Samples were weighed, homogenized in PBS and normalized to g/ml. To enumerate spore burdens, samples were heated at 70°C for 30 minutes, diluted and then plated onto *C. difficile* selection agar (containing cycloserine, cefoxitin, and fructose) supplemented with 0.1% w/v sodium taurocholate. After anaerobic incubation for up to 48 h viable counts were determined.

### Quantification of toxins in mice ceca

Cecal samples were collected and flash frozen. Samples (normalized to g/ml) were homogenized in PBS containing cOmplete^TM^ Protease Inhibitor Cocktail from Roche, then centrifuged at 10, 000 rpm for 5 min. The supernatants (5 µl) were used to quantify toxins by the above standard cell rounding assay.

## ACKNOWLEDGEMENTS

This work was funded by grants R56AI126881 and R01AI139261 to J.G.H. from the National Institute of Allergy and Infectious Diseases at the National Institutes of Health. The funders had no role in study design, in data collection and interpretation of the findings, or in the writing and submission of the manuscript.

