## Supplementary Tables S2-S3 and Figures S1-S3 for "The Ferrous Iron Transporter FeoB1 is Essential for *Clostridioides difficile* Toxin Production and Pathogenesis in Mice"

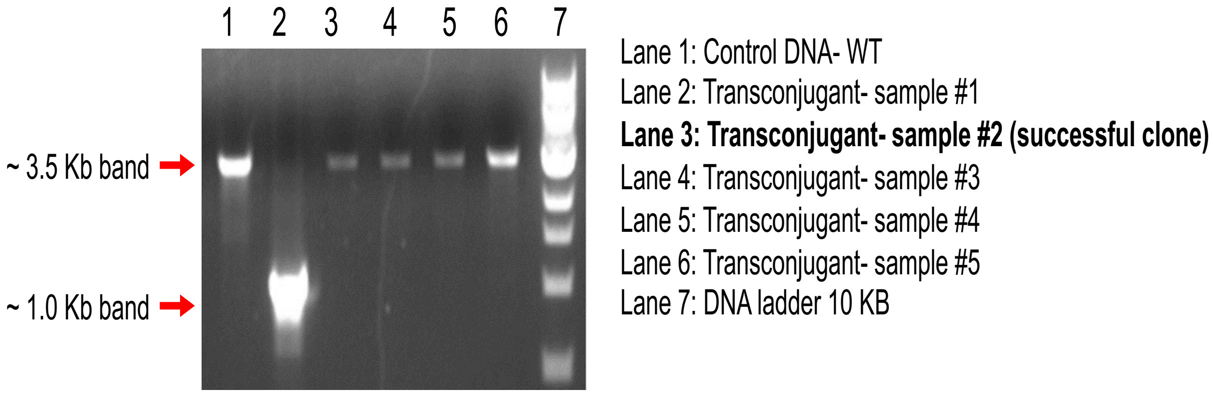


**Figure S1. PCR confirmation and screening of colonies lacking *feoB1*.** In above electrophoresis gel, transconjugants were purified on agars as being sensitive to thiamphenicol. DNA was extracted and PCR performed. Transconjugant sample 2 lacked *feoB1* as confirmed by ~2.5 kb reduction band size compared to the WT.

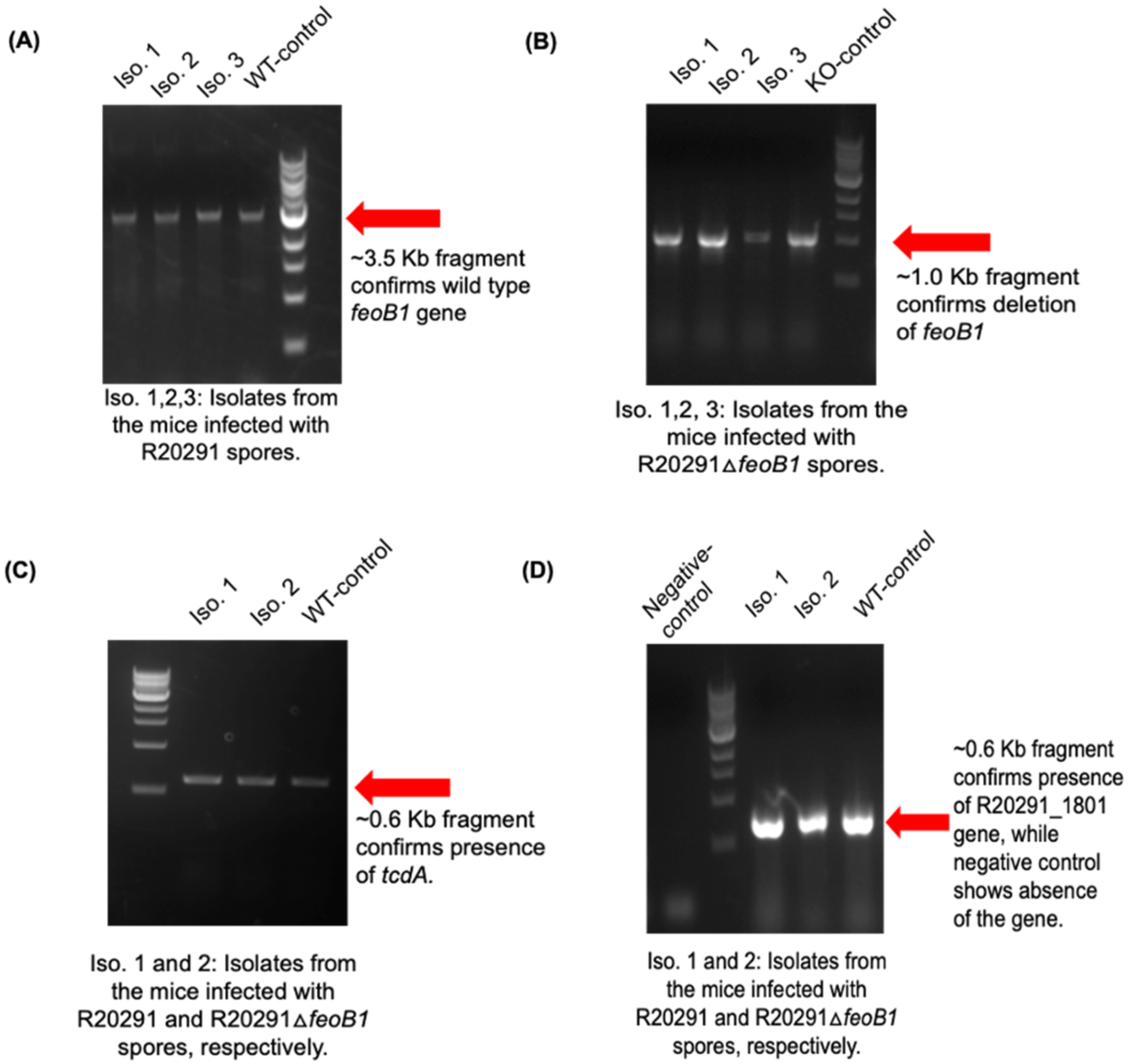


**Figure S2. PCR confirmation of the cecal isolates. A, B.** Random isolates were chosen for presence and absence of *feoB1* from cecal samples of mice infected with R20291 and R20291Δ*feoB1*, respectively. **C.** Toxin gene *tcdA* was confirmed in cecal isolates. **D.** Primers for gene *CDR20291_1801* which is found in ribotype 027 was confirmed in cecal isolates by PCR; CD196 was used as a negative control.

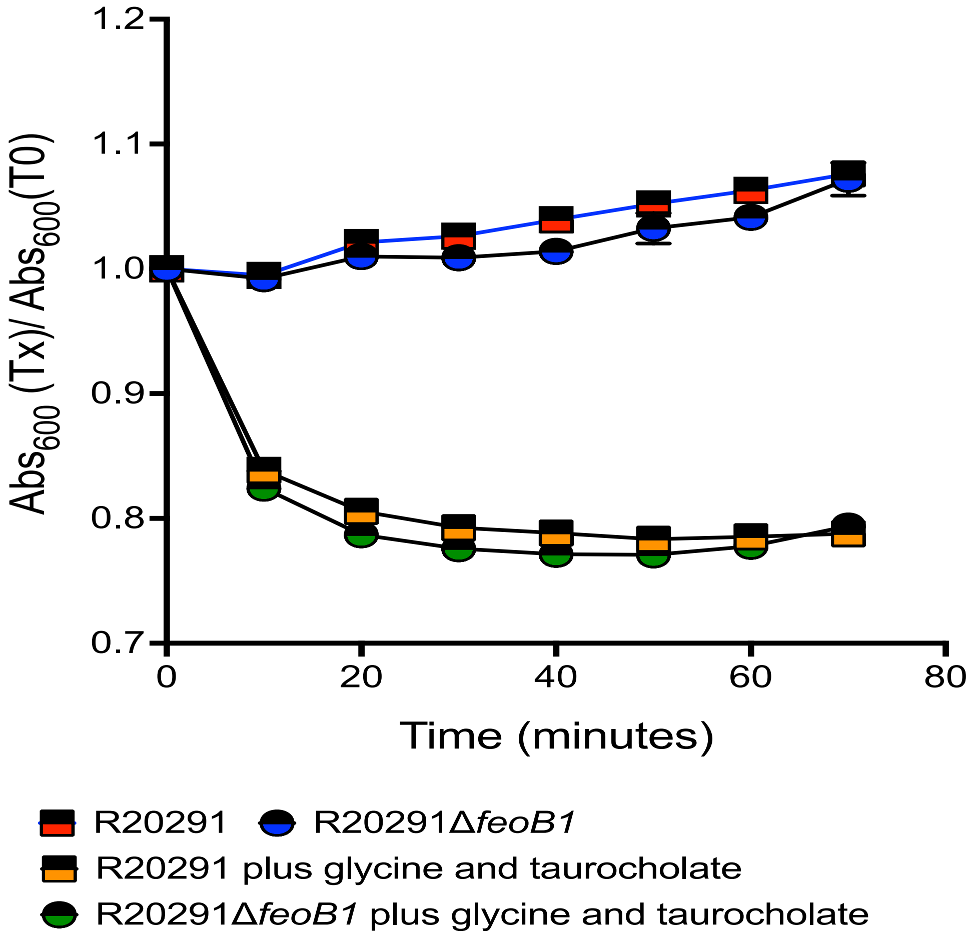


**Figure S3. Germination of R20291 versus R20291Δ*feoB1*.** Change in optical density based on time (Abs$_{600}$ (Tx) / Abs$_{600}$ (T0)) of spores in BHI in the presence or absence of the germinant taurocholate (0.1% w/v) and co-germinant glycine (10 mM).

**Table S2. Primers used in the study**

| Primer name | Description | Sequence |
|---|---|---|
| *feoB1*-F | To confirm deletion of *feoB1* | AGGTGAAGAAGTAGTTATAAAGAAAATTTCGGG |
| *feoB1*-R | To confirm deletion of *feoB1* | CTAATAATGCTCTTCCCAATCCACCGC |
| **For RT-qPCR experiments** | | |
| *fldx*_qPCR_FP1 | To analyze the expression of *fldx* | GATAAATGTTATCAATAAGTATTGAATTTCAAAATCAATTGTGATATTC |
| *fldx*_qPCR_RP1 | | CTCCCATAGATGGGCAACCTAATGC |
| *ribB*_qPCR_FP | To analyze the expression of *ribB* | CCGTAAATGGAGTTTGTCTAACTGTTTC |
| *ribB*_qPCR_RP | | GATTTATGGTTACTACTATTAGAATTTAATTCCTTGTAGTTATTAG |
| *ribD*_qPCR_FP | To analyze the expression of *ribD* | CAAAGGATATCATGAGAAATTTGGC |
| *ribD*_qPCR_RP | | GCAACATGTGAAAACTATAGTTTTACTGG |
| *bfr*_qPCR_FP (R20291_2756) | To analyze the expression of *bfr* | GAGTGATGCACATTACGACATGC |
| *bfr*_qPCR_RP (R20291_2756) | | CAATACTTCTCTATTGCCTTTTCAAATAAACTTAC |

| | | |
|---|---|---|
| *pyc*_qPCR_FP (R20291_0010) | To analyze the expression of *pyc* | GATAAGATTAACTCTAAAAAAATCGCAAAAGAAG |
| *pyc*_qPCR_RP (R20291_0010) | | GTTTTGGATCAGCTATATATTTTTCTATAAATATAATATCTTCTC |
| *acnB*_qPCR_FP (R20291_0763) | To analyze the expression of *acnB* | GGAGAAGCTGTTGCTGGAAGTTC |
| *acnB*_qPCR_RP (R20291_0763) | | CTCCATATTTATCTGCCACTGTTTGTATATACTTATG |
| *icd*_qPCR_FP (R20291_0764) | To analyze the expression of *icd* | GAAGTTGCGAAAGCTATGAAAAAGG |
| *icd*_qPCR_RP (R20291_0764) | | CTTAAATTGACATACAAATCTAATGCCTGTCTTAAAG |
| *fur_qPCR-FP* | To analyze the expression of *fur* | ACGCCACAAAGAAGAGCAAT |
| *fu*r_qPCR-RP | | TCAGGACAGTCAACCCTAACT |
| *fhuC*_qPCR-FP | To analyze the expression of *fhuC* | GGAAAATCGACAATACTTAAAACTATAGGTAGAATTATTAG |
| *fhuC*_qPCR-RP | | CAGTAACTTCTAAACTCCAATTTACAATCTCTC |
| *fhuD*_qPCR-FP | To analyze the expression of *fhuD* | GAGGTTAAGATACCATCAAATCCTAAAAAAATAG |
| *fhuD*_qPCR-RP | | GCTATTTCTTTTAATTGGTCATATATTTTTTCTTGTC |
| *tcdA*_qPCR-FP | | GGTGGAGAAGTCAGTGATATTG |

| | | |
|---|---|---|
| *tcdA*_qPCR-RP | To analyze the expression of *tcdA* | CAACTCCTGACTATAAATATTTAATAACTCTTG |
| *tcdB*_qPCR-FP | To analyze the expression of *tcdB* | GAAGAGTCATTAAATAAAGTTACAGAAAATAG |
| *tcdB*_qPCR-RP | | CGTTGTAGTATTGAAATCGTTATCC |
| *tcdD*_qPCR-FP | To analyze the expression of *tcdD* | GAAGAGGGAGAAACAGATTTAATAATATTTTTATTG |
| *tcdD*_qPCR-RP | | CCTCAAAAACAGACTTACTTTGATAATTATTTTTC |
| *tcdE*_qPCR-FP | To analyze the expression of *tcdE* | GGAGGCGTTATGAATATGACAATATC |
| *tcdE*_qPCR-RP | | GATGTTTTAGTCTTAAAAAATTGATACAATCTTG |
| IL-10_qPCR-FP | To analyze the expression of *il-10* | CGGGAAGACAATAACTGCACCC |
| IL-10_qPCR-RP | | CGGTTAGCAGTATGTTGTCCAGC |
| TNF-α _qPCR-FP | To analyze the expression of *tnf-α* | GGTGCCTATGTCTCAGCCTCTT |
| TNF-α _qPCR-RP | | GCCATAGAACTGATGAGAGGGAG |
| GM-CSF-qPCR_FP | To analyze the expression of *gm-csf* | AACCTCCTGGATGACATGCCTG |
| GM-CSF-qPCR_RP | | AAATTGCCCCGTAGACCCTGCT |
| β-actin-qPCR-FP | | CATTGCTGACAGGATGCAGAAGG |

| | | |
|---|---|---|
| | To analyze the expression of β-actin | |
| β-actin-qPCR-RP | | TGCTGGAAGGTGGACAGTGAGG |

**Table S3. Scoring chart criteria for animal health monitoring**

| Score | % Weight loss |
|---|---|
| 0 | normal appearance of fur and skin color; very active, alert, and excitable when observed |
| 1 | pink anogenital area; ruffled appearance of fur with normal skin color; moderate activity and alertness when observed |
| 2 | ruffled appearance of fur with normal skin color; little activity when observed (lethargic); wet tail |
| 3 | ruffled appearance of fur with normal skin color; little activity when observed (lethargic); wet tail |
| 4 | ruffled appearance of fur with abnormal skin color; wet tail; signs of paralysis, hunched posture, and a distended abdomen |
| **Score** | **General appearance** |
| 0 | < 5% |
| 1 | < 10% |
| 2 | < 15% |
| 3 | >15% - < 20% |
| 4 | >20 % |
| **Score** | **Temperature** |
| 0 | Normal from baseline (i.e., pre-infection) |
| 1 | Small changes of potential significance: 0.5°C < 1°C |
| 2 | Temperature change of 1 to < 3°C |
| 3 | Temperature change of ≥ 3°C |
| 4 | Temperature change of >20 % |
